# Flux Balance Analysis Identifies Distinct NADPH Production Strategies Across NCI 60 Cancer Cell Lines

**DOI:** 10.1101/332262

**Authors:** Zaid Ahmad

## Abstract

Flux Balance Analysis is a linear mathematical procedure which determines the set of reaction fluxes to produce a maximum flux of a reaction of interest. In this study, a core cancer model developed by Zielinski et al. 2017 is constrained by a set of 59 cancer cell type specific uptake and secretion rates. Optimizing for cell type specific biomass objective reactions and examining serine flux distributions reveals variability in production of NADPH. In many cell lines, production of NADPH is correlated to biosynthetic demand, however, outliers exist that produce excess NADPH beyond that of biomass demand. These outliers are first characterized by their NADPH production strategy (pentose phosphate pathway or a combination of One Folate Cycle and Malic Enzyme) and then the factors responsible for the different NADPH production strategies are identified. Results indicate that pentose phosphate pathway (PPP) producing NADPH cell lines had reprogrammed tricarboxylic acid cycle metabolism to meet the demand for decreased flux through glycolytic enzymes, while one folate cycle and malic enzyme (OFC + ME) producing NADPH cell lines had higher threonine, tyrosine and serine uptake.

## Introduction

Metabolic reprogramming has long been associated as a feature of developing cancer cells and tumorigenesis, however, recently there has been a revival of interest in the process by which cancer cells adapt to new cellular objectives of increased proliferation and maintain a tumorigenic state (Pavlova and Thompson, 2016). Although cancer cells have greater uptake of a variety of essential nutrients, studies have shown that cancer cells do not use this increased uptake to increase utilization of pathways fueled by these same nutrients in normal cells, specifically cancer cells do not, in general, have significantly increased production of ATP compared to normal cells (Pavlova and Thompson 2016). Instead, cancer cells aim to maximize biomass production for continuous proliferation and optimize NADPH to support biomass production (Pavlova and Thompson 2016). NADPH, in addition to functioning as a reducing agent in biomass producing reactions, also serves to prevent oxidative stress by reactive oxygen species (ROS) (Zielinski et al., 2017).

Although the pentose phosphate pathway has long been recognized as the major contributor to the production of NADPH, with malic enzyme also playing a role, the one carbon cycle downstream of the serine synthesis pathway has also, more recently, been shown to have an important role in contributing to NADPH production (Fan et al., 2014). However, flux through the serine synthesis pathway has been shown to be highly variable and while flux is elevated in some cancer cell lines, in other cancer cell lines, exogenous serine is the sole source of serine production (Mattiani et al., 2016). In addition to having in an important role in redox control, the SSP also has been shown to have strong implications with nucleotide biosynthesis (Mehrmohamadi et al., 2014). Studies have also shown that extracellular serine and glycine fails to support cell proliferation in the presence of inactivated SSP, providing further evidence for the role of the SSP beyond serine production (Mattiani et al., 2016).

In a previous study, Zielinski et al., determined that combined production of NADH and NADPH, as well as uptake of various amino acids such as serine, exceeded biomass demands and that these results were largely due to a high rate of glycolysis (Zielinski et al., 2017). Using the same core cancer model constrained by a set of 59 NCI 60 cancer cell biomass reactions, uptake/secretion rates, and demand reaction constraints, this study aimed to use flux balance analysis to determine the factors driving excess NADPH production based on the three common sources: PPP, OFC and malic enzyme reaction. As an in silico modeling technique that allows researchers to trace changes in individual reaction fluxes and link these to changes in different cellular goals or constraints, flux balance analysis is useful in making comparisons of the same model constrained differently (Ghaffari et al., 2015) (Orth and Palsson, 2010). Thus, differences in measured NADPH production can be linked to differences in constraints. Cytosolic NADPH was studied distinctly from mitochondrial NADPH or mitochondrial NADH in order characterize the contribution due to the various sources of NADPH from malic enzyme, pentose phosphate pathway and one folate cycle downstream of the serine synthesis pathway. The underlying purpose of the study was to determine how differences in unique nutrient profiles in cancer cell lines contribute to distinct NADPH production strategies.

## Methods

### Setup

A core cancer model developed by Zielinski et al 2017 was downloaded from the supplementary information of the article at doi:10.1038/srep41241. In addition to the core cancer model, the measured uptake rates for 24 metabolites, along with the predicted biomass flux constraint and the ATP sink constraint (defines the maximum and minimum ATP production predicted based on measured values) was downloaded from the supplementary excel file. 59 cell specific biomass reactions, created based on the biomass compositions of the in vitro cells, came with the supplementary models. (Zielinski et al., 2017).

Matlab R2016b, Cobra Toolbox 2.0, and Gurobi 7.5.1 were used to carry out the Flux Balance Analysis in various conditions. The main cobra toolbox functions that were used were optimizeCbModel.m, changeRxnBounds.m, and changeObjective.m.

### Protocol

#### Contribution of Serine Uptake and Serine Synthesis Pathway to Total Serine Production

The biomass optimization data was used to determine the relative contribution of various sources of serine production. First the total production and consumption of cytoplasmic serine was determined by taking all the reactions that produced serine and adding the fluxes and taking all the reactions that consumed serine and adding their fluxes. As expected, due to the inherent mass-charge balance, serine production equaled serine consumption. Serine consumption that was to be allocated for biomass purposes was determined by multiplying the serine stoichiometric coefficient in the biomass reaction by the overall biomass reaction flux. The percent contribution of serine production from exogenous sources for each of the 59 cell line models was calculated and plotted against various reactions relating to serine metabolism in the pathway.

#### Calculation of Sources of NADPH Production

Cytosolic NADPH production for each cell line was measured by combining the fluxes of methylene tetrahydrofolate dehydrogenase, formyl tetrahydrofolate dehydrogenase, malic enzyme and glucose 6 phosphate dehydrogenase. The total NADPH production was plotted against biomass reaction flux and was characterized based on the contribution of serine synthesis pathway reactions that produced cytosolic NADPH, the contribution of malic enzyme to total cytosolic NADPH production, and the contribution of pentose phosphate pathway flux to total NADPH production.

Mann Whitney two tailed u tests were performed using matlab ranksum.m function. Each sample consisted of one set of reactions for three possible conditions: High cytosolic NADPH production resulting from high malic enzyme activity and high flux through one folate reactions downstream of serine synthesis pathway, high cytosolic NADPH due to pentose phosphate pathway and low cytosolic NADPH that was correlated with biomass reaction flux.

## Results

### Characterization of the Serine Synthesis Pathway

Percent contribution of serine from exogenous sources was shown to be highly variable across the 59 modified cancer models and ranged from 14.09 percent for the CCRF-CCM leukemia cell line model to 100 percent for multiple cell lines; as there were only two sources of serine production-the SSP and exogenous serine, percent contribution of serine from exogenous sources was inverse to utilization of the SSP (Figure 1). Comparison of percent contribution of serine from exogenous sources to serine uptake flux, comparing the percent contribution of serine from exogenous sources to the flux of the phosphoserine phosphatase reaction (that catalyzes the conversion of 3-phosphoserine to serine), greater variability in the phosphoserine phosphatase reaction was found at lower percent contributions than at higher ones. An identical trend was found when percent contribution of serine from exogenous sources was compared to overall serine production – that is percent contribution ranges of lower values were shown to have greater variability than percent contribution ranges of higher values (Figure 1). Additionally, all reactions of the one folate carbon cycle were correlated to the production of serine through phosphoserine phosphatase, except those constrained by other sources (thymidylate synthetase, dihydrofolate reductase and aminoimidazole carboxamide formyltransferase) (Figure 2).

**Figure 1:**
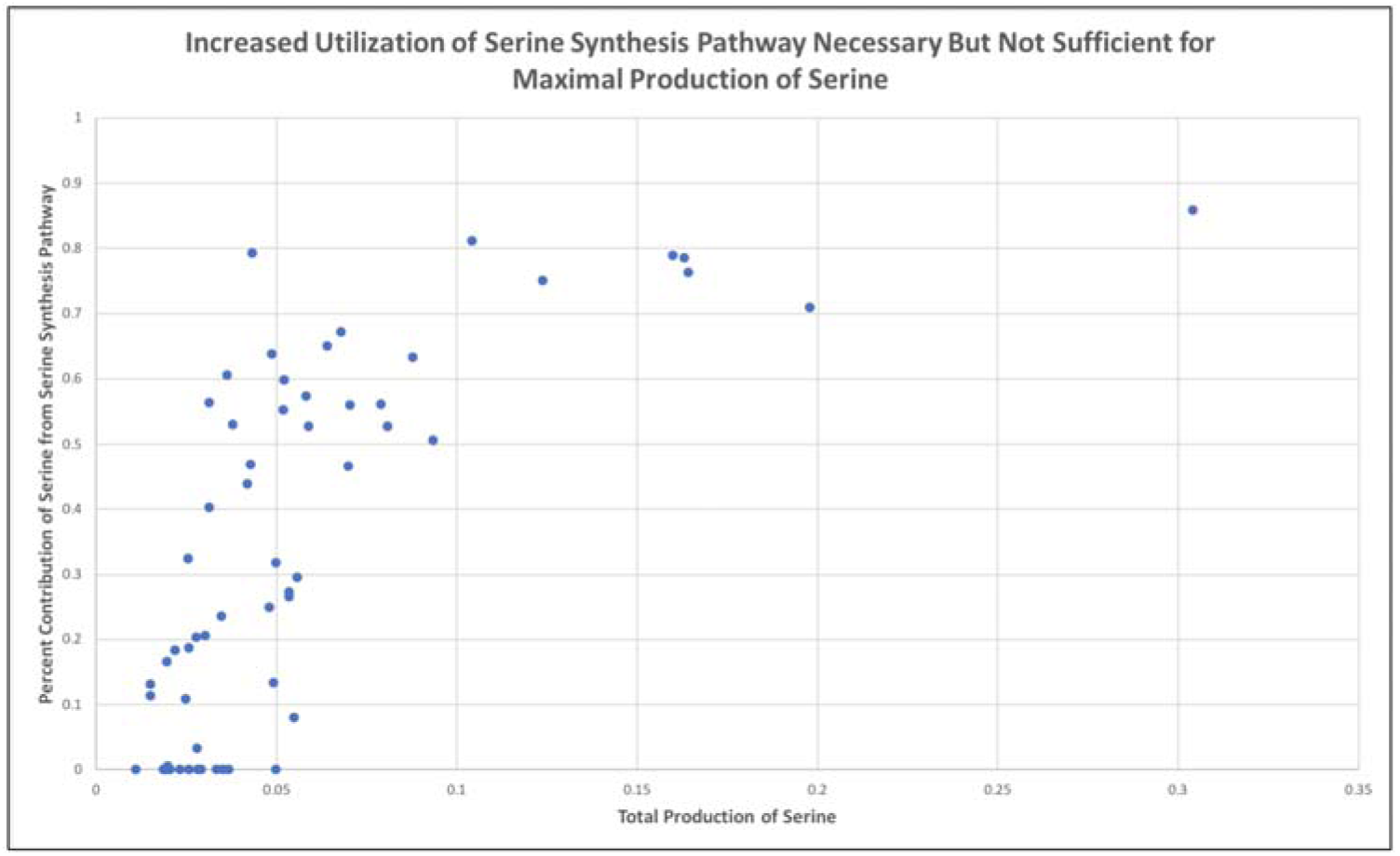
Total production of serine is highly variable and reaches its maximum capacity at higher utilizations of the phosphoserine phosphatase reaction to produce serine. At higher utilizations of the serine synthesis pathway, maximal serine production can be reached by pairing of phosphoserine phosphatase serine production with serine uptake from exogenous sources.

**Figure 2:**
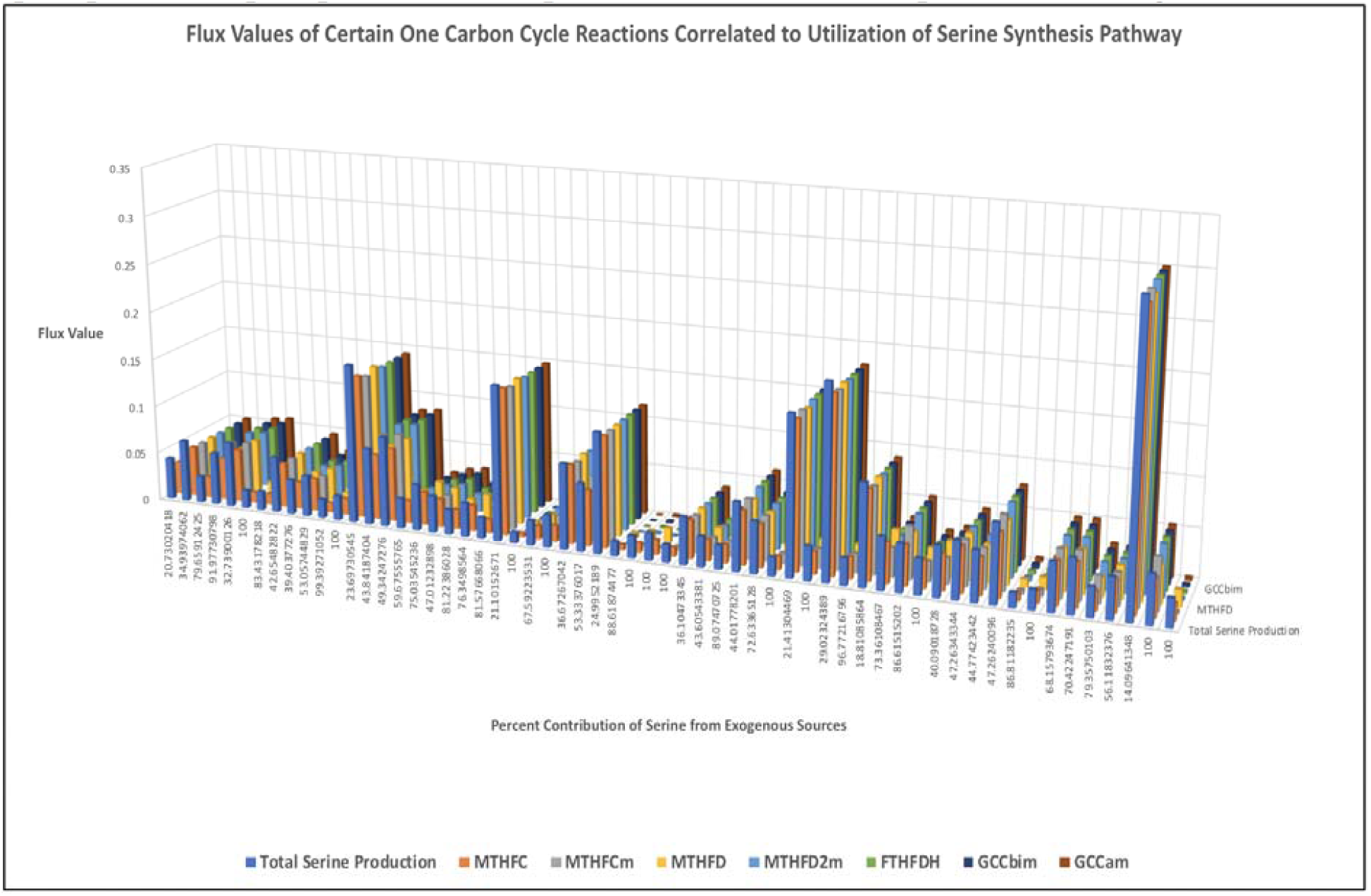
Many, but not all, reactions involved in One Folate Cycle are correlated with utilization of the Serine Synthesis Pathway. Reactions not correlated are constrained by other factors.

### Total Production of Cytosolic NADPH

Total production of cytosolic NADPH was calculated by adding together the flux values of all cytosolic NADPH producing reactions. Total production of cytosolic NADPH was plotted against each of the three cytosolic reactions that consumed NADPH as well as to biomass reaction flux (Figure 3). A trend of increased NADPH correlated to increased biomass reaction flux was observed across all cell lines except for 13 outliers. These outliers included 3 colon cancers (2 from PPP condition, 1 from OFC + ME condition), 1 breast cancer (PPP), 2 brain cancer (1 from PPP; 1 from OFC + ME), 3 ovarian cancers (1 from PPP and 2 from OFC + ME), 2 leukimia (both OFC + ME), 1 renal cancer (OFC + ME) and 1 prostate cancer (OFC + ME). High NADPH producing cell lines either had high PPP flux/low SSP flux/low malic enzyme flux or had low PPP flux and a combination of high malic enzyme flux, high utilization of serine synthesis pathway and high total production of serine, thus demonstrating pairing of high serine uptake with utilization of SSP and ME to produce NADPH comparable to that produced only from PPP. Cell lines with lower NADPH (NADPH that directed biomass reaction flux and was thus a reaction flux determining factor) had zero or significantly lower pentose phosphate pathway flux than high NADPH from PPP and had either high SSP or high serine uptake rate, producing lower production of SSP (Figure 4).

**Figure 3:**
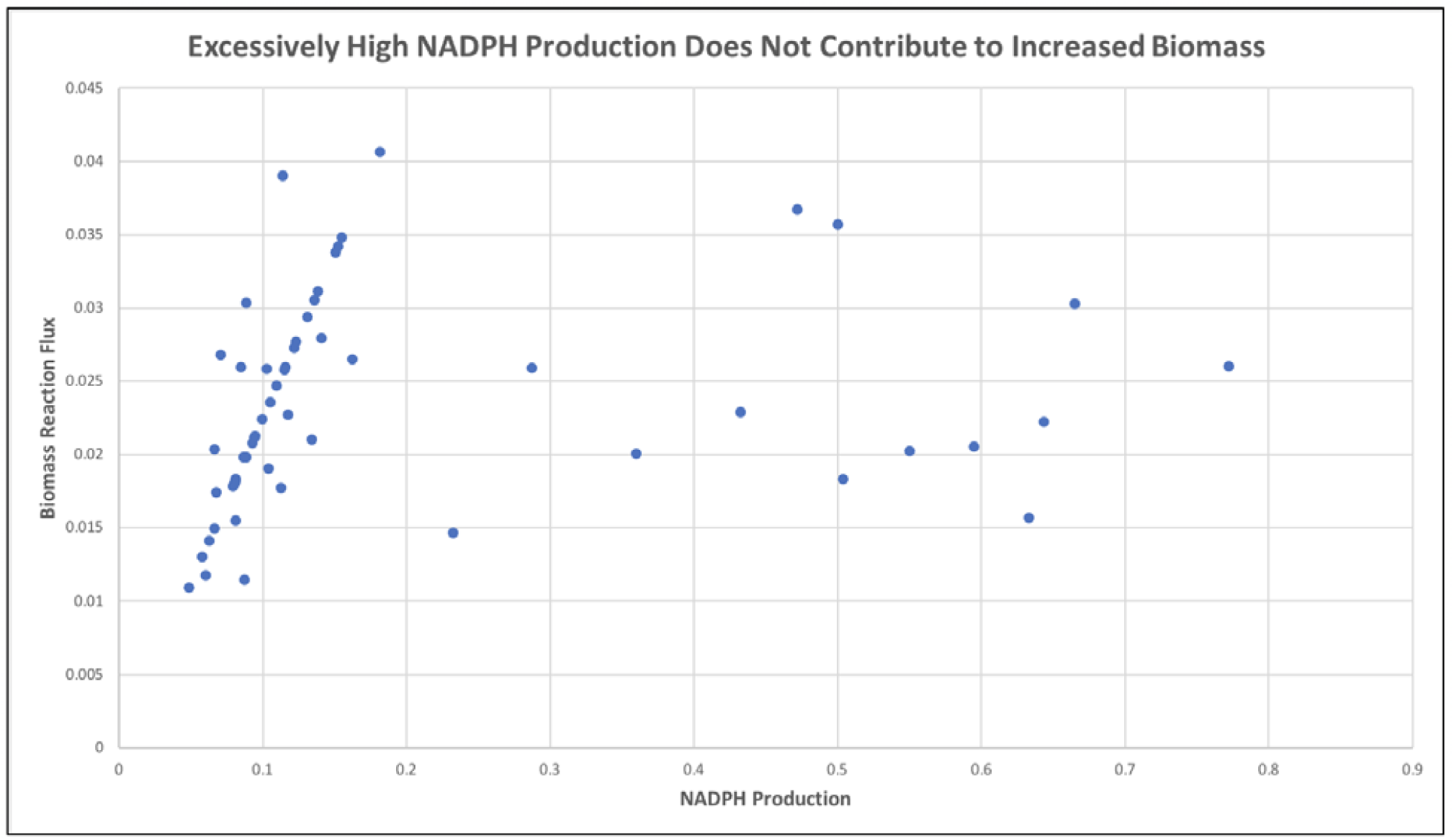
Above a certain threshold, high NADPH production does not contribute to increased biomass reaction flux. High NADPH production can result from high flux through the one folate cycle in combination with malic enzyme or through high flux through the pentose phosphate pathway.

**Figure 4:**
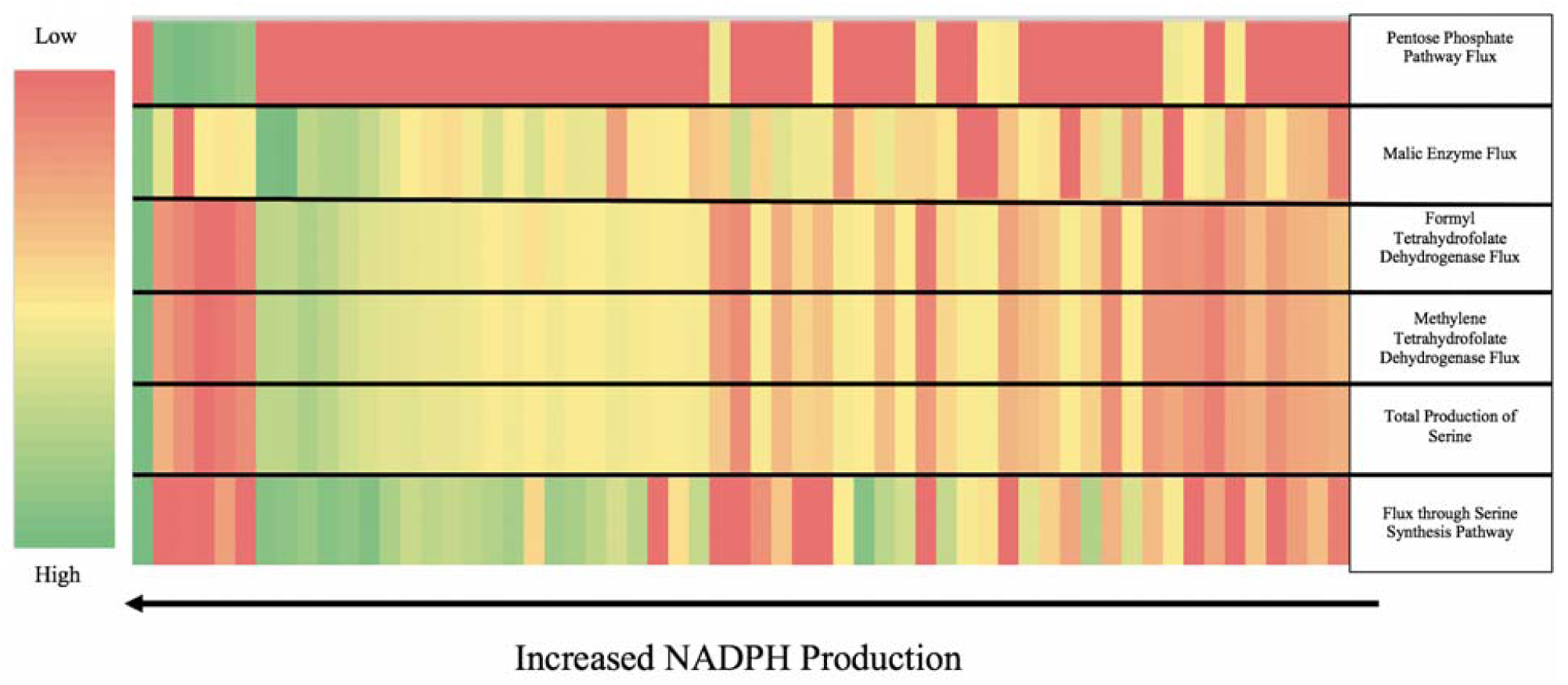
As total NADPH production increases, cells utilize different sources to supply NADPH. Flux through serine synthesis pathway gradually increases as does total production of serine and one folate reactions supplying NADPH to increase total production of serine. At higher total production of NADPH, either malic enzyme pairs with one folate reactions driven by high serine production to increase overall NADPH or, at even higher concentrations pentose phosphate pathway supplies nearly all of the NADPH.

### Statistical Analysis

Statistical Analysis of the two high NADPH producing conditions revealed that serine uptake, threonine uptake and tyrosine uptake were all significantly greater in the High NADPH from OFC + ME condition compared to the High NADPH from PPP condition (Figure 5). Statistical analysis of flux variability, which finds the maximum and minimum possible bounds for each reaction, demonstrated that the maximum possible flux through glycine intake into the mitochondria was significantly greater in the Low NADPH and High NADPH from OFC + ME condition compared to the High NADPH from PPP, further providing evidence for the fact that serine production serves to meet glycine biomass demand. Reactions that demonstrated significant differences between the High NADPH from OFC + ME and High NADPH from PPP included SSP reactions, one folate cycle reactions and reactions involving amino-oxo butane. Reactions that were not a part of this pathway but did show significant differences included the aspartate glutamate mitochondrial shuttle (PPP favored mitochondrial aspartate production while OFC + ME favored mitochondrial glutamate production), chloride and glutamate transport (transport into the mitochondria was less in both cases for PPP) and the bicarbonate equilibration reaction (Figure 6) (Supplementary Results). These results provide evidence of reprogrammed tricarboxylic acid cycle metabolism to compensate for decreased flux through glycolysis.

**Figure 5:**
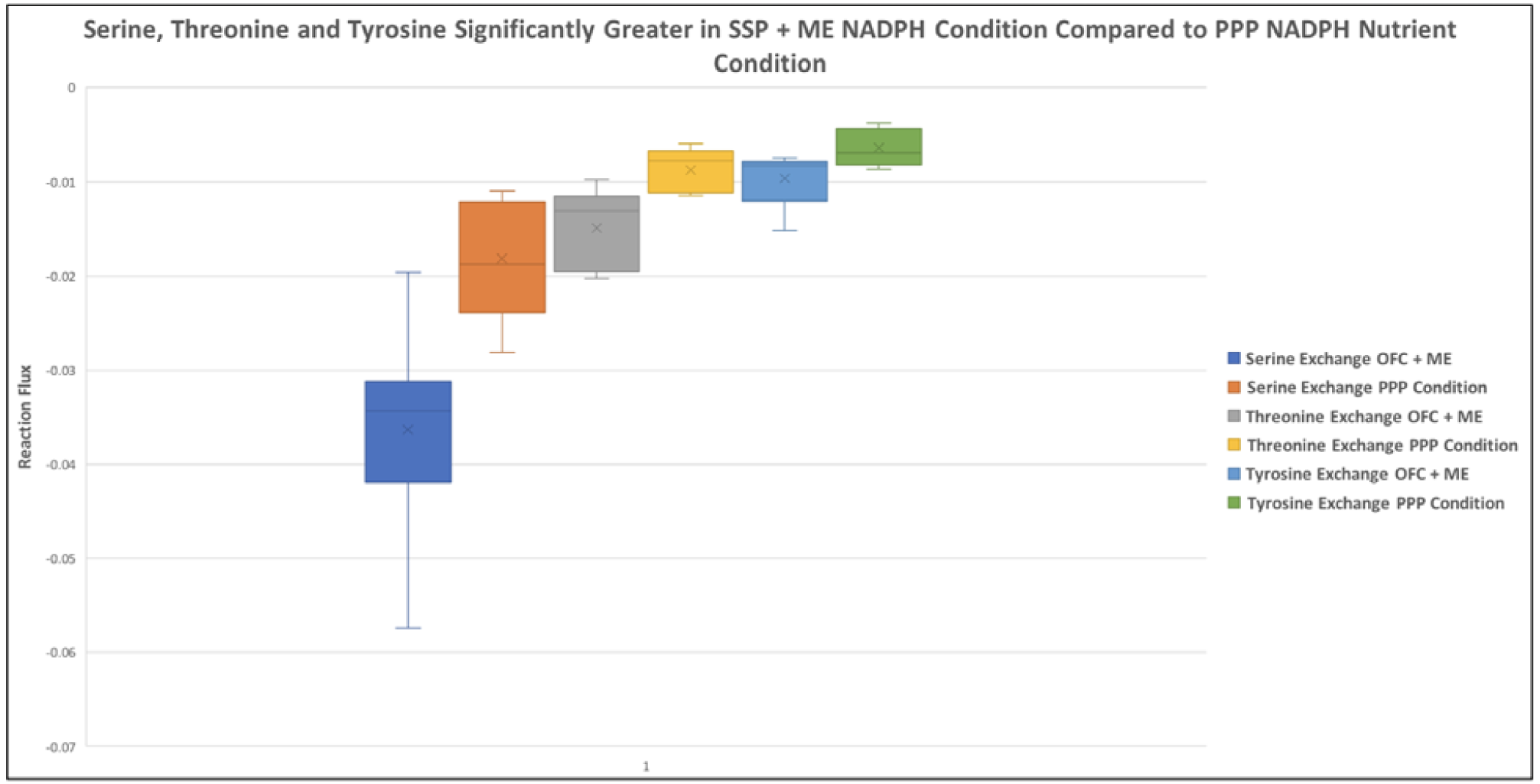
Threonine, tyrosine and serine are increased in OFC + ME high NADPH condition as compared to PPP high NADPH condition. Threonine uptake allows for glycine production thus allowing serine to be utilized for increased flux through one folate cycle.

**Figure 6:**
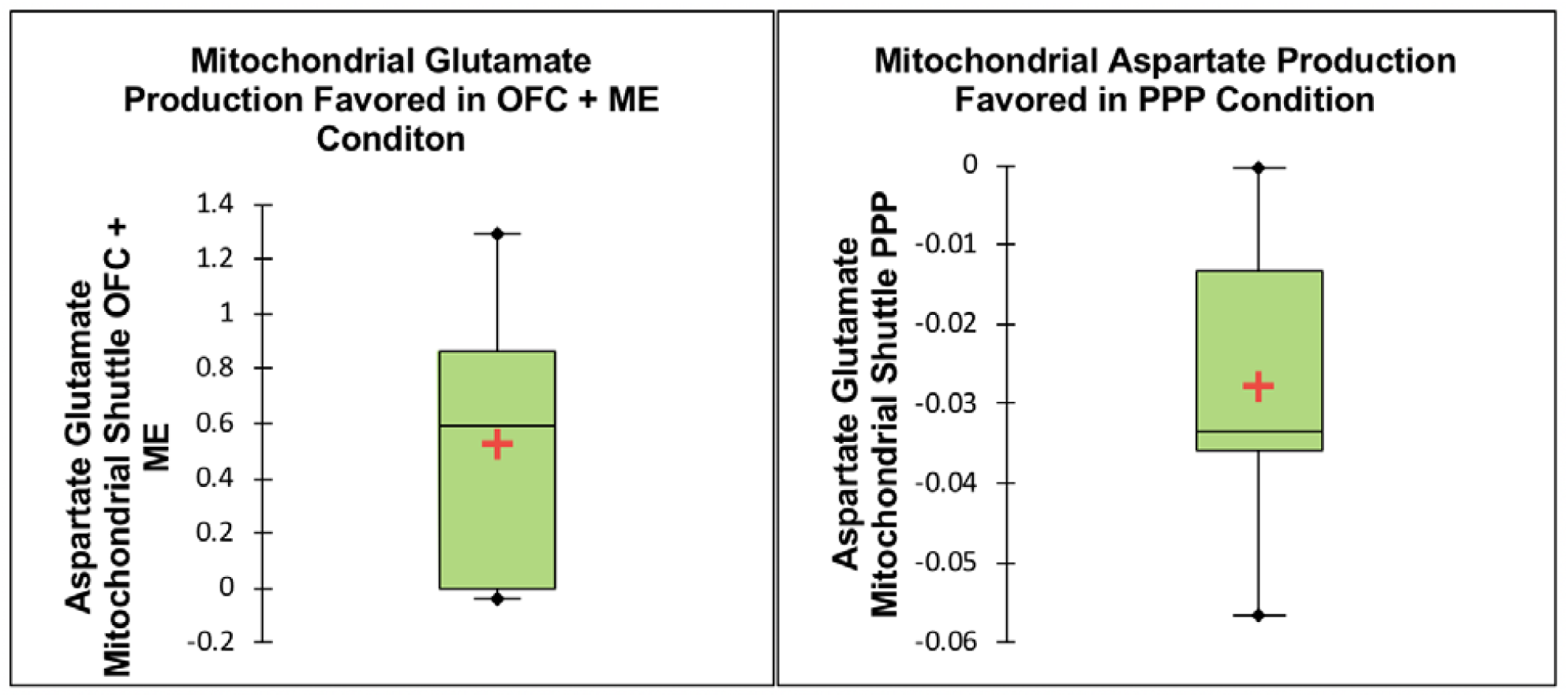
Flux through aspartate glutamate shuttle indicative of mitochondrial metabolic reprogramming in high NADPH producing cells. Mitochondrial aspartate produced in PPP condition can be used for production of mitochondrial oxaloacetate, possibly compensating for decreased production of oxaloacetate by mitochondrial pyruvate.

Statistical comparison of the High NADPH from PPP condition compared to statistically significantly lower NADPH was carried out to determine the factors contributing to high Pentose Phosphate Pathway flux. In addition to flux through glycolytic reactions downstream of the pentose phosphate pathway shunt being significantly less in the PPP condition, reactions of the Tricarboxylic acid cycle were also less. Production of mitochondrial alpha ketoglutarate was favored in malate alpha ketoglutarate shuttle compared to production of mitochondrial malate being favored in the malate alpha ketoglutarate shuttle in the low NADPH condition (Figure 7) (Supplementary Results). In the mitochondrial aspartate transaminase reaction, production of mitochondrial oxaloacetate was favored in the PPP condition. Mitochondrial glutamate from this reaction was then converted to mitochondrial aspartate. Similar to the OFC + ME condition, the HCO3 equilibration reaction flux was also less in the PPP condition as compared to the low NADPH condition.

**Figure 7:**
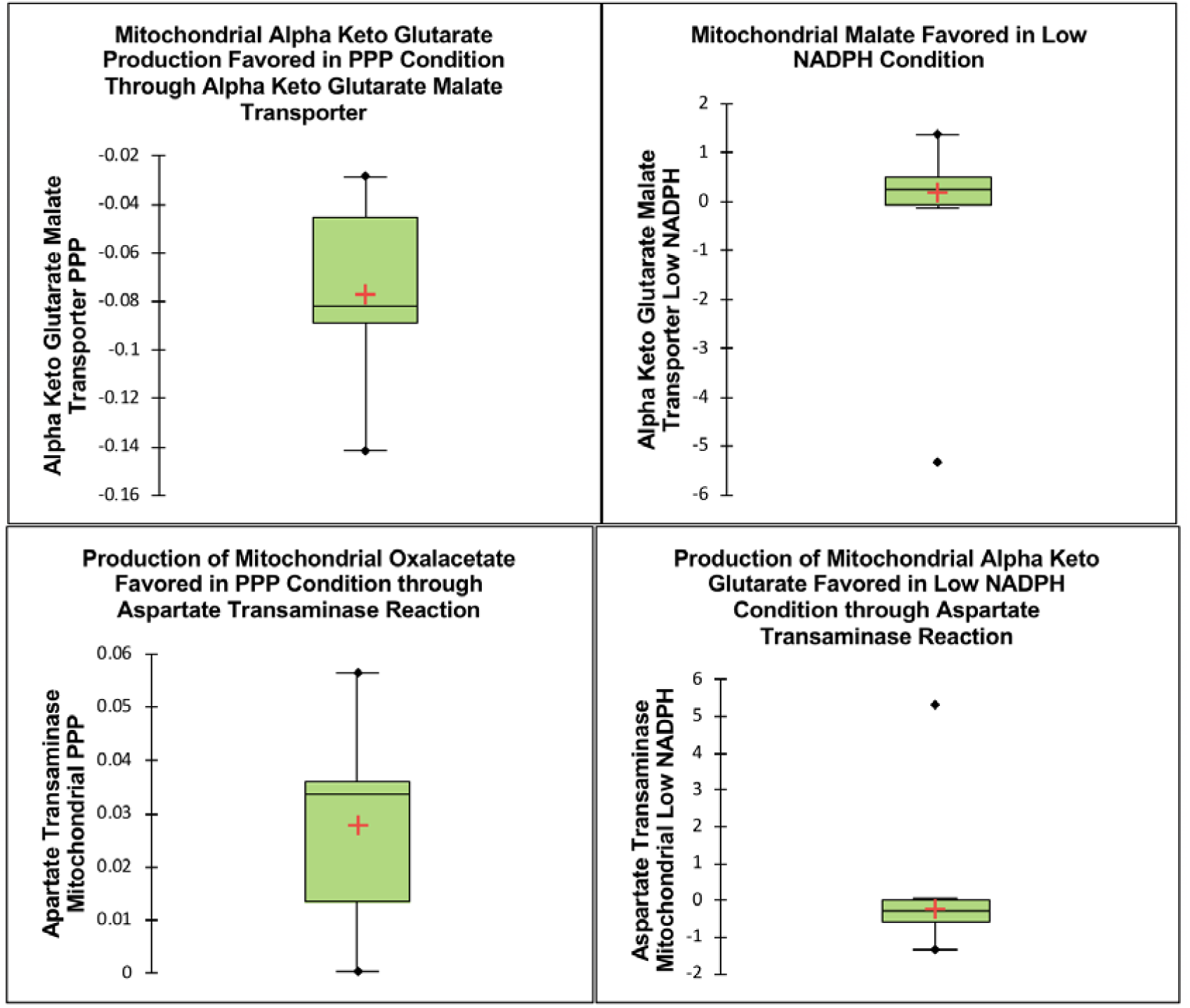
In the PPP condition, mitochondrial alpha keto glutarate is favored in the malate alpha keto glutarate transporter, allowing for production of mitochondrial oxaloacetate in the aspartate transaminase reaction. In the low NADPH condition, mitochondrial alpha keto glutarate is made by the aspartate transaminase reaction. Mitochondrial alpha keto glutarate produced from the malate alpha keto glutarate shuttle in the PPP condition is used to produce oxaloacetate, thus compensating for decreased oxaloacetate production from mitochondrial pyruvate.

## Discussion

As the various routes to NADPH production have been characterized in the past, this study sought primarily to characterize various NADPH production strategies in cancer cell lines and secondarily, determine the factors that contribute to excess NADPH production in the context of cancer. Although serine uptake was shown to play an important role in the total production of serine, excess serine production that was not driven by biomass demand was only achieved by pairing serine uptake with malic enzyme flux and high utilization of the serine synthesis pathway. Alternatively, in certain cell lines pentose phosphate pathway was able to produce high levels of NADPH on its own and without the need of other contributors of NADPH. As certain cancers such as ovarian and colon cancers had some cell lines that produced high NADPH from the PPP and some that produced high NADPH from the OFC + ME, it was hypothesized that constraining serine uptake would lead to a switch to utilization of the pentose phosphate pathway to produce NADPH. However, constraining serine uptake instead led to significantly lower levels of NADPH, thus implying that factors contributing to increased NADPH production through the pentose phosphate pathway were distinct from factors contributing to increased NADPH production through the SSP and ME.

The fact that threonine uptake was significantly greater in the OFC + ME condition compared to the PPP condition may provide a clue as to how SSP flux is controlled and mediated. Threonine can be converted to amino-oxo butane which can enter the mitochondria and produce mitochondrial glycine. In a previous study using the same models used in this study, Zielinski et al. determined that serine uptake may serve to meet the biosynthetic demands of both serine and glycine. In cell lines with higher uptake of threonine, threonine may provide a source for production of glycine, thus allowing serine produced through the serine synthesis pathway and serine taken in from exogenous sources to be used for production of NADPH and increased flux through the one folate cycle. In contrast, although in the OFC + ME condition, glycolysis was not significantly decreased, in the PPP condition, glycolytic reactions and tricarboxylic acid reactions saw a significant decrease. This is due to the fact that 2 intermediates of 3 phosphoglycerate are produced for each one molecule of glucose, however this is not true for glucose 6 phosphate. Additionally, reprogrammed transport of the mitochondrial-cytosolic transport mechanisms, with a focus on increased production of oxaloacetate may serve as a mechanism for meeting biosynthetic demand as well as producing high amounts of cytosolic NADPH.

## Conclusion

A central strategy in targeting cancer metabolism is to target the metabolic differences between cancer cells and normal cells (Selwan et al. 2016). Previously studies have shown the distinct differences between nutrient profiles of various types of cancer (Jain et al., 2012). Understanding the effects of uptake of various nutrients in cancer cell lines is useful due to the fact that altered redox balance is one of the hallmarks of cancer cells and is responsible for drug resistance in many cancers (Panieri and Santoro 2016). Additionally, characterizing the cancer specific alterations that allow for variable NADPH production may prove useful in developing therapeutic strategies targeted for different cancer cell lines. The fact that constraining uptake of nutrients such as serine in high NADPH cell lines leads to low NADPH production implies that cancer cell lines develop specific strategies for NADPH production based on their specific nutrient profiles. Selectively altering or inhibiting nutrient uptake may thus lead to constrained NADPH production while also allowing for decreased biomass, however, these strategies should be cancer cell line specific.

## Acknowledgements

I would like to thank Dr. Daniel Zielinski at University of California San Diego and Dr. Larryn Peterson at Rhodes College Departament of Chemistry for his help in navigating this project.

